# Evolution of myoglobin in avian lineages: positive selection for aggregation resistance in penguins and high purifying selection in galliforms

**DOI:** 10.1101/2021.02.26.433090

**Authors:** Syed M. Rizvi, Wei Zheng, Chengxin Zhang, Yang Zhang

**Affiliations:** Department of Computational Medicine and Bioinformatics, University of Michigan, Ann Arbor, MI 48109 USA; Department of Biological Chemistry, University of Michigan, Ann Arbor, MI 48109 USA

## Abstract

Myoglobin is the major oxygen carrying protein of vertebrate muscle, and high myoglobin net charge is known to hold evolutionary significance as a molecular signature of secondarily aquatic diving capacity in mammals. However, the evolution of myoglobin’s electrostatic properties in non-mammalian vertebrates, such as birds, has not been investigated. Here, we used a new deep learning-based protein folding algorithm to model the tertiary structures of myoglobin from 302 vertebrate species and performed a comparative analysis of their net charge, positively charged solvent-accessible surface area, and negatively charged solvent-accessible surface area. For avian myoglobins, we also calculated selection pressure (ω). The results suggest that the myoglobins of diving avians, specifically those of the penguins (*Sphenisciformes*) and diving ducks (*Aythyini*), have highly positively charged electrostatic surfaces, which evolved via positive selection to reduce aggregation propensity and allow greater storage of oxygen for extended underwater foraging. In contrast, galliform myoglobins are under high purifying selection. Distribution of charged atoms on myoglobin surface was more indicative of high myoglobin content than net charge. We also found inter-class differences in net charge; bird myoglobins are the most positively charged and reptile and amphibian myoglobins are the most negatively charged, and net charge seems to be negatively associated with herbivory within mammals. Finally, we propose an equation that describes the relationship between myoglobin net charge and concentration better than the previously suggested logarithmic function. Our findings offer novel insights into the diversification of myoglobin in vertebrate clades and highlight the power of computational structural approaches for zoological and evolutionary research.

## Introduction

Myoglobin, the metalloprotein responsible for red color and oxygen storage in muscle tissues, is a model protein for understanding structure-function relationships.^1,2^ Myoglobin holds special significance for molecular and evolutionary physiologists due to its adaptive diversification in various mammalian clades.^3–6^ For example, diving mammals cannot breathe underwater, so they store the oxygen acquired from surface breathing into tightly packed myoglobin reserves for use during extended dives.^6,7^ Myoglobin of such mammals tends to have a high net charge, creating electrostatic repulsion between myoglobin proteins to prevent the deleterious aggregation expected at such high intracellular myoglobin concentrations. The discovery of this molecular signature helped position myoglobin as a model protein for understanding the evolution of secondarily aquatic mammals at a molecular level.^6^ Because protein aggregation is implicated in several degenerative diseases^8,9^ and also complicates the development of soluble biopharmaceuticals,^10,11^ the ability of myoglobins from secondarily aquatic mammals to avoid aggregation at the dangerously high concentrations found in these species is of special interest.^7^

Although myoglobin is one of the most well-studied proteins, the evolution of extreme myoglobin properties in non-mammalian vertebrate groups, such as birds, has not been investigated. For example, it is not known whether the myoglobins of diving birds (e.g. penguins) show adaptations to aquatic lifestyles,^6^ and information on variation in selective pressure across avian myoglobin lineages is also unavailable. Due to lack of studies on non-mammalian myoglobins, it is unknown whether there are any inter-class differences in vertebrate myoglobin net charge, and whether the differences have functional significance. Apart from other vertebrate groups, even within mammals the exact molecular nature of the relationship between myoglobin’s net charge and its intracellular concentration levels is not known.^6^ We also wonder if electrostatic properties other than net charge are more relevant for functional myoglobin electrostatic repulsion, especially tertiary structure-based properties such as the percent of solvent-accessible surface area covered by positively charged atoms, which previous studies did not investigate.

To address these questions, we used a new deep learning-based protein folding algorithm to model the tertiary structures of myoglobin from 302 vertebrate species and performed a comparative analysis of their net charge, positively charged solvent-accessible surface area, and negatively charged solvent-accessible surface area. For avian myoglobins, we also calculated selection pressure (ω). Furthermore, we reexamined the relationship between myoglobin net charge and intracellular concentration levels with a larger dataset than previous studies and identified a better equation to describe the data along with biological and physical explanations for the equation’s terms.

## Methods

### Tertiary structure modeling by D-I-TASSER

To calculate structural properties (e.g. individual atoms’ contributions to solvent-accessible surface area) and accurately predict myoglobin net charge using the three-dimensional structure-based net charge prediction algorithm PROPKA3,^12^ D-I-TASSER^13^ was used to first create tertiary structure models of the 311 vertebrate myoglobin orthologs on the NCBI Orthologs website (https://www.ncbi.nlm.nih.gov/gene/4151/ortholog/?scope=7776) and the 95 additional mammalian species in the supplementary materials of Mirceta et al. 2013. As an extended pipeline of I-TASSER^14^, the D-I-TASSER^15^ protocol integrates deep convolutional neural network-based distance and hydrogen-bonding network prediction to guide the assembly of template fragments into full-length model by Replica-Exchange Monte Carlo (REMC) simulations. The pipeline consists of four consecutive steps. First, D-I-TASSER uses DeepMSA^16^ to iteratively search the query protein sequence against the whole-genome and metagenome sequence databases to obtain a multiple sequence alignment (MSA). Next, the selected MSA is used as the input for DeepPotential,^17^ a newly developed deep residual neural-network-based predictor, in order to create multiple spatial restraints including (1) distance-maps for both C*α* and C*β* atoms; (2) C*α*-based hydrogen-bonding networks; (3) C*α*-C*β* torsion angles. Meanwhile, LOMETS3, a newly developed meta-server program containing both profile- and contact-based threading programs, is used to identify structural templates from a non-redundant PDB structural library. In the third step, continuous fragments extracted from the template structures by LOMETS3 are assembled into the full-length decoys by a REMC simulation under the guidance of a composite force field consisting of the deep learning-predicted spatial restraints, template-derived distance restraints, and knowledge-based energy terms calculated based on statistics of PDB structures. Finally, decoys from low-temperature replicas generated by the REMC simulations are clustered by SPICKER.^18^ The SPICKER clusters are refined at the atomic level using fragment-guided molecular dynamic (FG-MD) simulations,^19^ with the side-chain rotamer structures repacked by FASPR.^20^

### Estimation of structure model quality

The global quality of structural models is usually appraised by the TM-score^21^ between modeled and experimental structures:

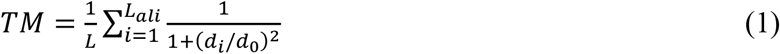

where *L* is the number of residues, *d*_*i*_ is the distance between the *i-*th aligned residue pair, and 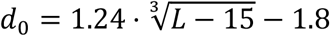 is a scaling factor. TM-score ranges between 0 and 1, and a TM-score greater than 0.5 indicates a structure model of correct global topology.^22^

In the present study, experimental structures were not available for 99% of the myoglobin orthologs. Therefore, instead of the actual TM-score, we estimate the TM-score (*eTM-score*) using threading alignment quality, contact map fulfillment rate, and simulation convergence in D-I-TASSER:

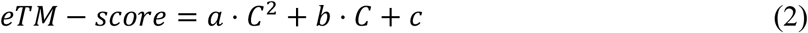

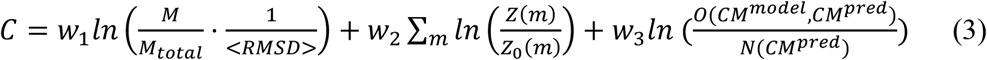

where *a* = 0.00098, *b* = 0.10770, and *c*= 0.79 are free parameters retrieved by regression, *C* is the c-score, *M*_*total*_ is the total number of decoy conformations used for clustering, *M* is the number of decoys in the top cluster, <*RMSD*> is the average RMSD of decoys in the same cluster, *Z*(*m*) is the score of the top template by threading method *m, Z*_*0*_(*m*) is the cutoff for reliable templates, *N*(*CM*^*pred*^) is the number of predicted contacts that guide the REMC simulation, and *O*(*CM*^*model*^, *CM*^*pred*^) is the number of overlapping contacts between final models and predicted contacts. *w*_1_ = 0.77, *w*_2_ = 1.36 and *w*_3_ = 0.67 are free parameters. Estimated TM-score (*eTM-score*) highly correlates with real TM-score, with Pearson Correlation Coefficient (PCC) 0.81 on 797 training proteins.^23^

### Data selection

To ensure accuracy of the final models, six models with estimated TM-score^21^ <0.5 were removed from the analysis. Following strict statistics of structures in the PDB, TM-scores higher than 0.5 assume generally the same fold.^22^ Unusually short orthologs (< 140 residues) were also discarded. Because some of the models included long N-terminal tails which resulted in unrealistically high net charge, STRIDE^24^ was used to assign the N-terminal coil/loop regions for all models, and these regions were manually removed. After these curations, the final analysis included a total of 392 vertebrate myoglobins, of which 224 were from mammals, 73 from birds, 72 from fish, 21 from reptiles, and 2 from amphibians.

### Electrostatics analyses

Net charge of the folded state for each of the myoglobin models was calculated using the PROPKA3 algorithm,^12^ under the physiologically relevant pH of 7. Visualization of surface electrostatics (figure 5) was performed using the APBS Electrostatics plugin. PyMOL scripts (supplementary text S1) were designed to calculate and visualize solvent-accessible surface areas.^25^

### Selection pressure (ω) and phylogenetic history

Selection pressures (ω), or the dN/dS ratios, were calculated in MATLAB with Needleman–Wunsch alignment. Evolutionary relationships between the bird orders in this study (figure 3) were extracted from an established comprehensive phylogeny construction of birds based on 259 nuclear loci and >390,000 bases of genomic sequence data from each of 198 avian species, highly supported by bayesian and maximum likelihood analyses.^26^

### Diet

Diet information for species was retrieved using web searches and confirmed with the MammalDIET database.^27^

### [Mb] values and curve fitting

Maximal skeletal muscle myoglobin concentrations of 31 mammalian species were retrieved from the supplementary materials of Mirceta et al. for testing associations between concentration, and net charge.^7^ Curve fitting was performed using the curve fitting tool in MATLAB.

## Results and Discussion

### D-I-TASSER model quality

The global quality of structural models can be assessed by the TM-score^21^ between modeled and experimental structures of the target protein. TM-score ranges between 0 and 1, with TM-score > 0.5 meaning structure models of correct global topology.^22^ At the time of this writing, experimentally determined structures for four of the vertebrate species in our dataset were available in the Protein Data Bank (PDB): *Homo sapiens* (PDB ID: **<u>3RGK</u>**), *Equus caballus* (PDB ID: **<u>1WLA</u>**), *Phoca vitulina* (PDB ID: **<u>1MBS</u>**), and *Physeter catodon* (PDB ID: **<u>1VXH</u>**). TM-scores between our predicted models and the experimentally determined experimental structures were greater than 0.8 for all four species (figure 1B). As the experimental structures for the 298 remaining orthologs are not available, we estimated the TM-score (*eTM-score*) of the D-I-TASSER models using a combination of threading alignment quality, contact satisfaction rate, and convergence of the structure assembly simulations (see details in Materials and Methods).^28^ In the current study, the estimated TM-score was greater than 0.8 for 94.4% of the 302 myoglobin models (figure 1B), and the estimated TM-score was close to the actual TM-score for the four species with experimentally determined structures (figure 1A).

**Figure 1.**
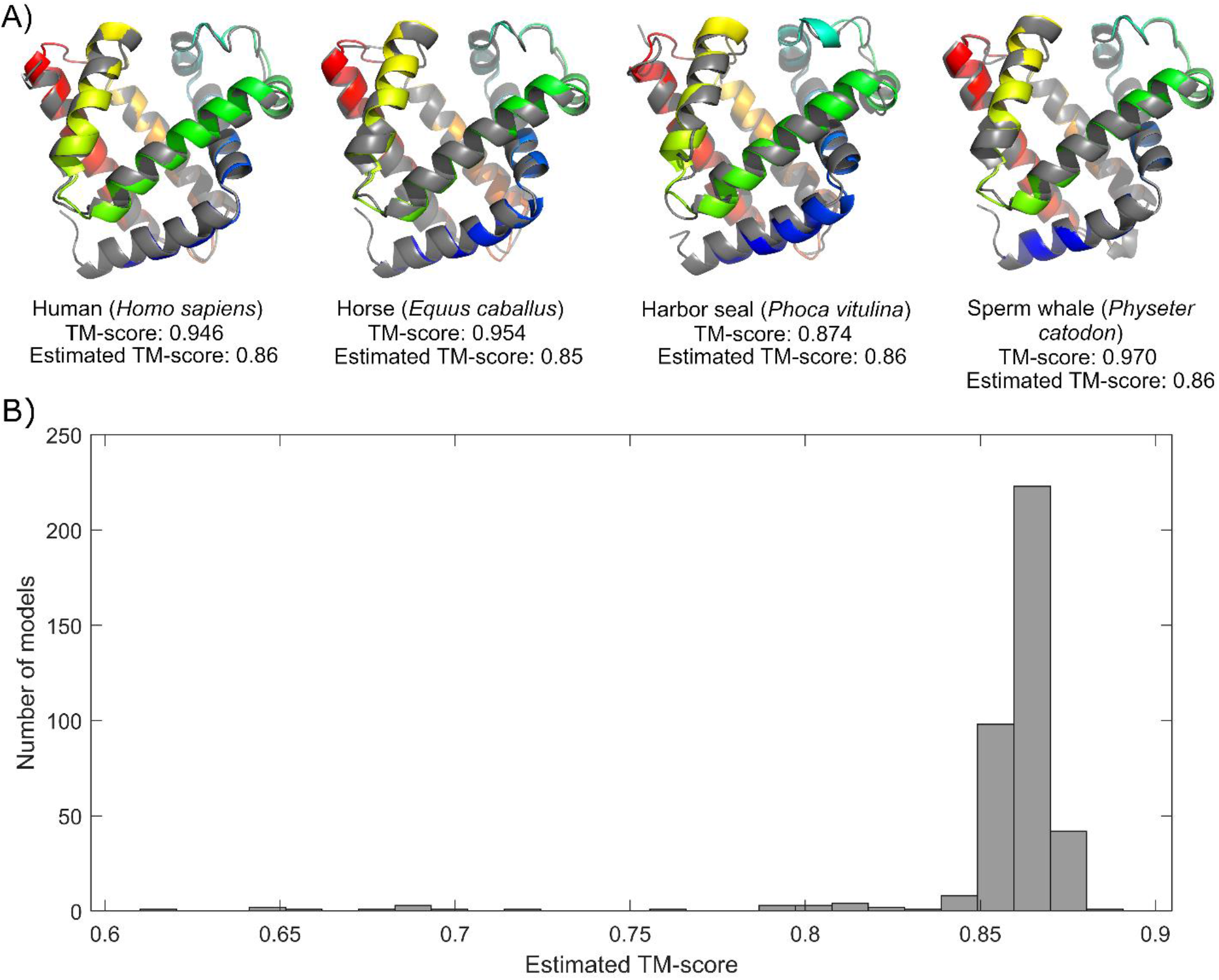
**(A)** TM-align superposition of models and experimental structures from human, horse, harbor seal, and sperm whale. **(B)** Histogram depicting the distribution of estimated TM-scores for the 302 D-I-TASSER models used in this study.

### Confirmation of positive correlation between logarithm of experimentally determined muscle myoglobin content and computationally modelled net charge

To test the validity of our methods, we tried to reproduce the findings of previous studies by examining the relationship between the logarithm of experimentally determined muscle myoglobin concentration (log[Mb]) data from previous literature and our predicted net charge for 69 of the mammalian species in our dataset for which measured myoglobin concentrations existed in previous literature.^7^ A positive Pearson correlation coefficient of 0.7468 (p-value = 1.73×10^−13^) was found between the predicted net charge and log[Mb], which suggests a strong positive linear correlation between experimentally measured data and our computationally modeled net charge (figure 2). This reproduces previous findings and indicates the role of increased electrostatic repulsion in preventing aggregation at high intracellular myoglobin concentrations.^6,7^

**Figure 2.**
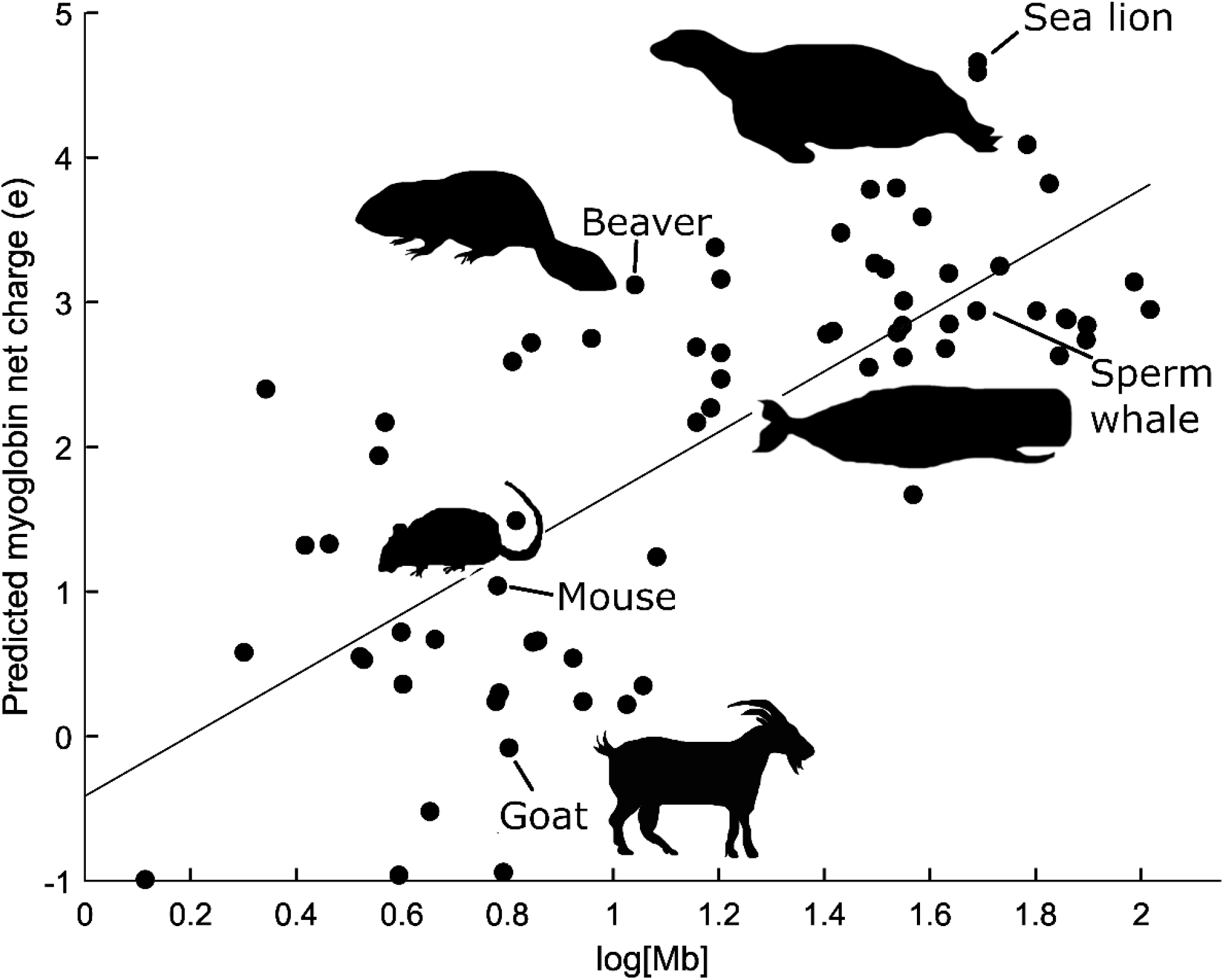
Predicted myoglobin net charge and logarithm of experimentally determined muscle myoglobin concentrations in mammals.

### High and low myoglobin content species differ more in tertiary structural features than net charge

Previous studies examined only one electrostatic property: net charge.^6^ However, given that the role of high net charge in myoglobin is to prevent aggregation via electrostatic repulsion, and that tertiary structure is more directly related to function than primary structure, we hypothesized that tertiary structural features that contribute to electrostatic repulsion might provide more information about myoglobin function and environmental pressures than net charge alone. In particular, we analyzed interspecific variation in the percent contribution of positively charged atoms (i.e. nitrogen atoms in the side chains of lysine, histidine, and arginine) and negatively charged atoms (i.e. oxygen atoms in the side chains of aspartic acid and glutamic acid) to the solvent-accessible surface area of myoglobin. We performed a correlation test and two-sample t-tests to determine whether high (90^th^ percentile) and low (10^th^ percentile) concentration species differ more in their positively charged solvent-accessible surface area than their net charge. The results partially confirmed our hypothesis; although the PCC between log[Mb] and predicted net charge (PCC = 0.7468, p-value = 1.73×10^−13^) was higher than the PCC between log[Mb] and percent positively charged solvent-accessible surface area (PCC = 0.6589, p-value = 7.50×10^−10^), the p-value for differences in mean percent positively charged surface area (p-value = 9.65×10^−7^) was lower than the p-value for differences in mean net charge (p-value = 2.41×10^−4^). These results suggest that increasing the percent positively charged solvent-accessible surface area is physiologically more important for protein aggregation resistance than net charge. This may have important implications for medicine; creating aggregation-resistant proteins is a crucial and unresolved problem for development of soluble biopharmaceuticals^10,11^ and redesign of the human proteins implicated in aggregation diseases (e.g. the design of aggregation-resistant crystallins to prevent cataracts, or aggregation-resistant amyloid precursors to prevent Alzheimer’s).^8,9^

### Penguins and diving ducks possess highly charged myoglobins adapted for aquatic hypoxia

Our avian dataset included five diving species and 68 non-diving species. The five diving birds are emperor penguin (*Aptenodytes forsteri*), Adélie penguin (*Pygoscelis adeliae*), tufted duck (*Aythya fuligula*), great cormorant (*Phalacrocorax carbo*), and red-throated loon (*Gavia stellata*). Significant differences in myoglobin net charge were found between diving and non-diving birds. Specifically, two major diving bird lineages, the *Sphenisciformes* (penguins) and *Anseriformes* (waterfowl), had higher average myoglobin net charge than the other bird orders (figure 3). Wilcoxon rank sum test revealed a statistically significant difference (p-value = 0.04) between the higher median net charge of the five diving species in our dataset (+3.55*e*) and the lower median net charge of the non-diving species (+2.11*e*). Diving birds in the 90^th^ percentile of net charge included the Emperor penguin (+3.57*e*), Adélie penguin (+3.55*e*), and the tufted duck (+6.56*e*), which is a species of diving duck. The tufted duck had the highest net charge of all birds (figure 4A). Comparison of surface electrostatics between diving and non-diving bird myoglobin models revealed obvious differences in charge intensity and localization, especially in the region dorsal to the heme binding site (figure 5).

**Figure 3.**
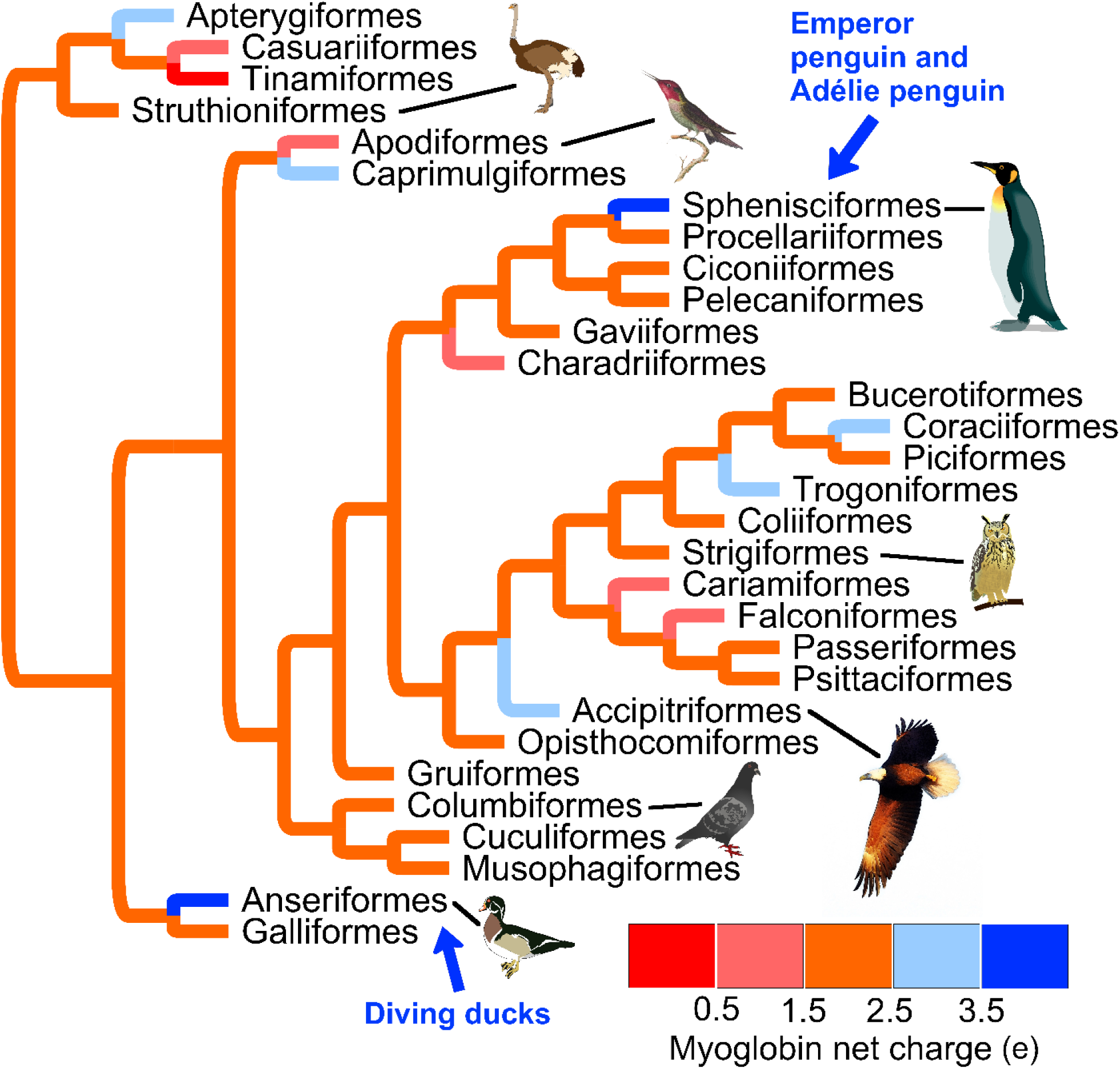
Cladogram and average predicted myoglobin net charge of bird orders. Branches between internal nodes were colored orange because the net charge range for 56.7% of bird orders falls between 1.5*e* and 2.5*e*.

**Figure 4.**
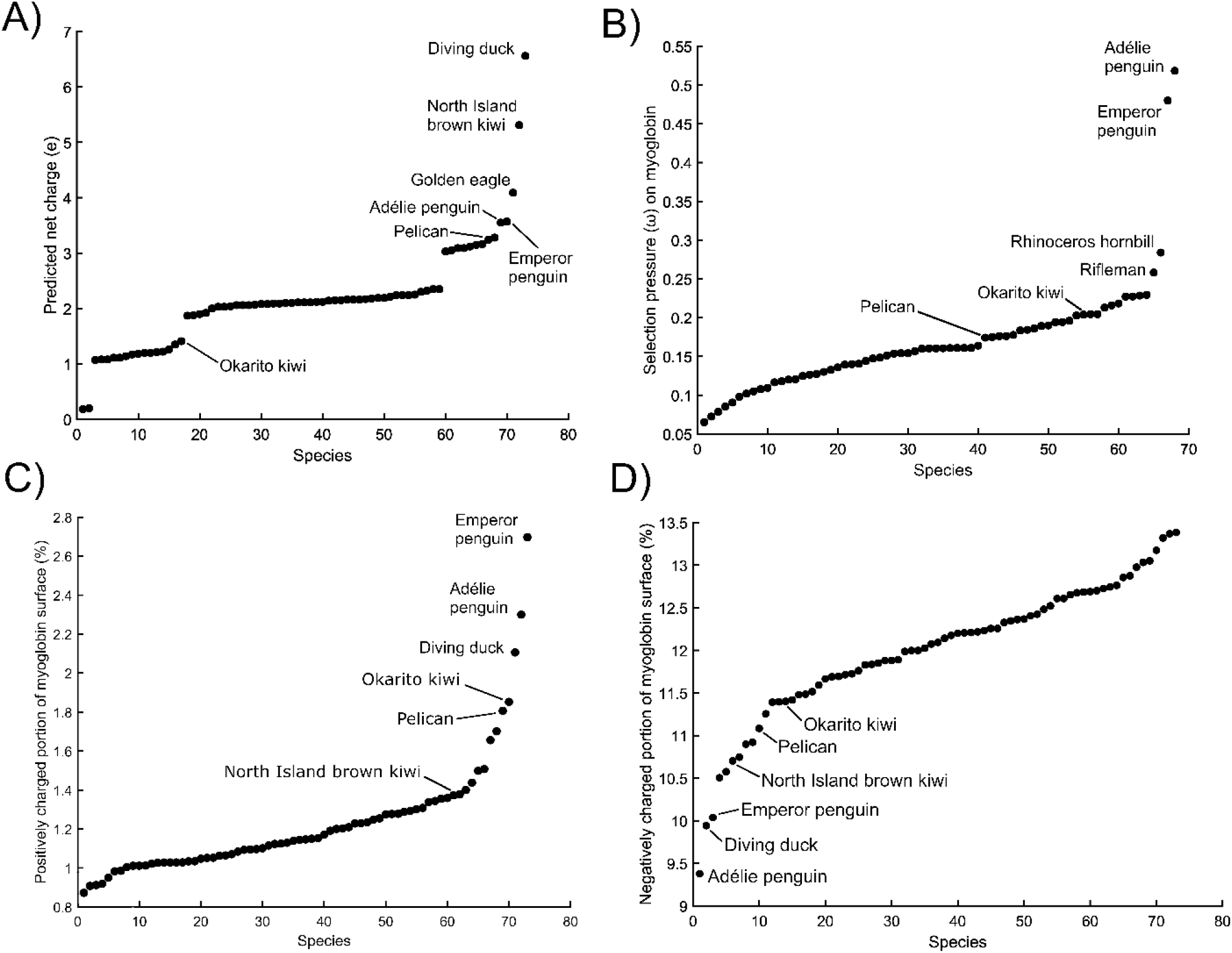
Scatterplots of **(A)** predicted myoglobin net charge, **(B)** myoglobin dN/dS ratio, or selection pressure (ω) on myoglobin, **(C)** contribution of positively charged atoms to myoglobin solvent-accessible surface area, and **(D)** contribution of negatively charged atoms to myoglobin solvent-accessible surface area, for myoglobin models of 73 bird species, each ranked from low to high in the respective measures.

**Figure 5.**
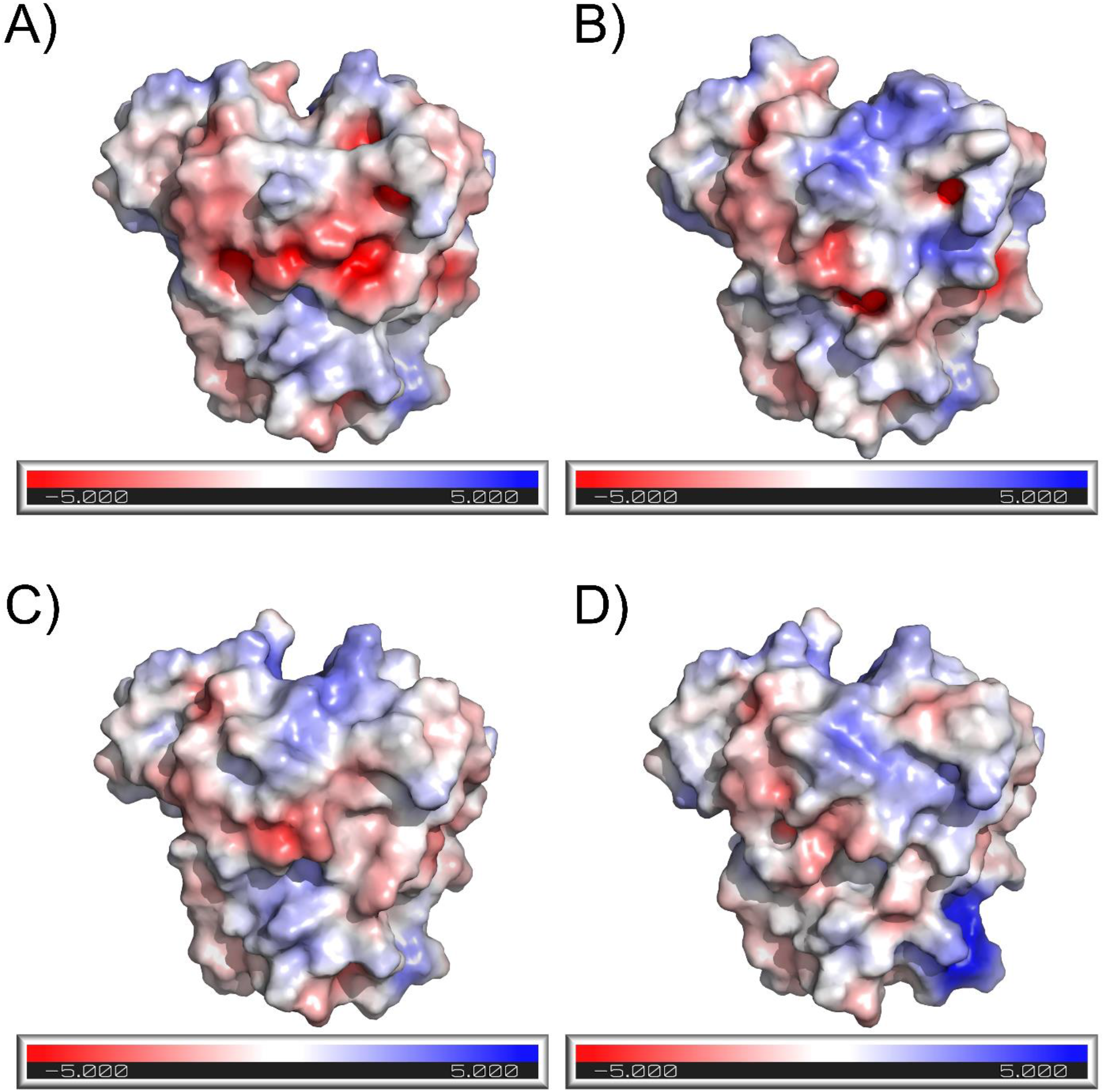
Visualization of surface electrostatics for the myoglobin models from **(A)** chicken, **(B)** Emperor penguin, **(C)** mallard, and **(D)** tufted duck. The side shown is dorsal to the heme binding site.

The link between myoglobin net charge and diving capacity depends on the assumption that the muscles of diving species possess unusually high concentrations of myoglobin. Although information on bird muscle myoglobin content is scarce, previous studies confirmed that three of the species in the 90^th^ percentile of myoglobin net charge in our study indeed possess unusually high muscle myoglobin content: tufted duck, Emperor penguin, and Adélie penguin.^29–31^ Penguins are known to possess one of the highest myoglobin concentrations in the entire vertebrate clade (64 mg/g in Emperor penguins),^29^ and the pectoral muscle myoglobin concentration of non-diving Muscovy ducks is 2.2 mg/g,^32^ which is significantly lower than the 7.4 mg/g for dive-trained diving ducks.^30^ Although penguins and diving ducks are known to have higher myoglobin content, it was unknown whether they also possess highly charged myoglobins or whether they evolved an alternative mechanism to address the increased likelihood of myoglobin aggregation.^6^ The associations we found between aquatic bird diving capacity, muscle myoglobin concentration, and myoglobin net charge solve this problem and suggest that mammalian and avian divers convergently evolved high net charge as a defense against myoglobin aggregation.

Structural electrostatic properties of penguin and diving duck myoglobins were even more pronounced than their net charge. Wilcoxon rank sum test revealed a statistically more significant difference between the median percent positively charged solvent-accessible surface areas of diving and non-diving birds (p-value = 0.0033) than the difference between their median net charges (p-value = 0.04), suggesting that myoglobin’s solvent-accessible surface electrostatics are a better predictor of organism niche than net charge. Whereas the net charge of the North Island brown kiwi (*Apteryx mantelli*) and golden eagle (*Aquila chrysaetos*) was higher than the net charge of penguins, the penguins and diving duck had the highest positively charged solvent-accessible surface areas and the lowest negatively charged solvent-accessible surface areas out of all the birds in our dataset (figure 4). These structural measures distinguish the electrostatic properties of penguins and diving ducks from all other avian species. As seen in Figure 6, a visual comparison of the contribution of positively charged atoms to solvent-accessible surface area reveals a greater contribution in the Emperor penguin than the ostrich (*Struthio camelus*). Furthermore, as seen in Figure 6, the contribution of negatively charged atoms to solvent-accessible surface area reveals a more attenuated contribution in the diving-adapted tufted duck than the closely related non-diving mallard (*Anas platyrhynchos*). The high percent of positively charged solvent-accessible surface area may provide stronger electrostatic repulsion and a higher likelihood of repulsion when myoglobins contact each other under stochastic conditions in the cytoplasm of myocytes. Because natural selection increased repulsive properties using positive-positive repulsion instead of negative-negative repulsion, attenuating the amount of negatively charged solvent-accessible surface area would reduce electrostatic attraction between the proteins and hence decrease aggregation propensity.

**Figure 6.**
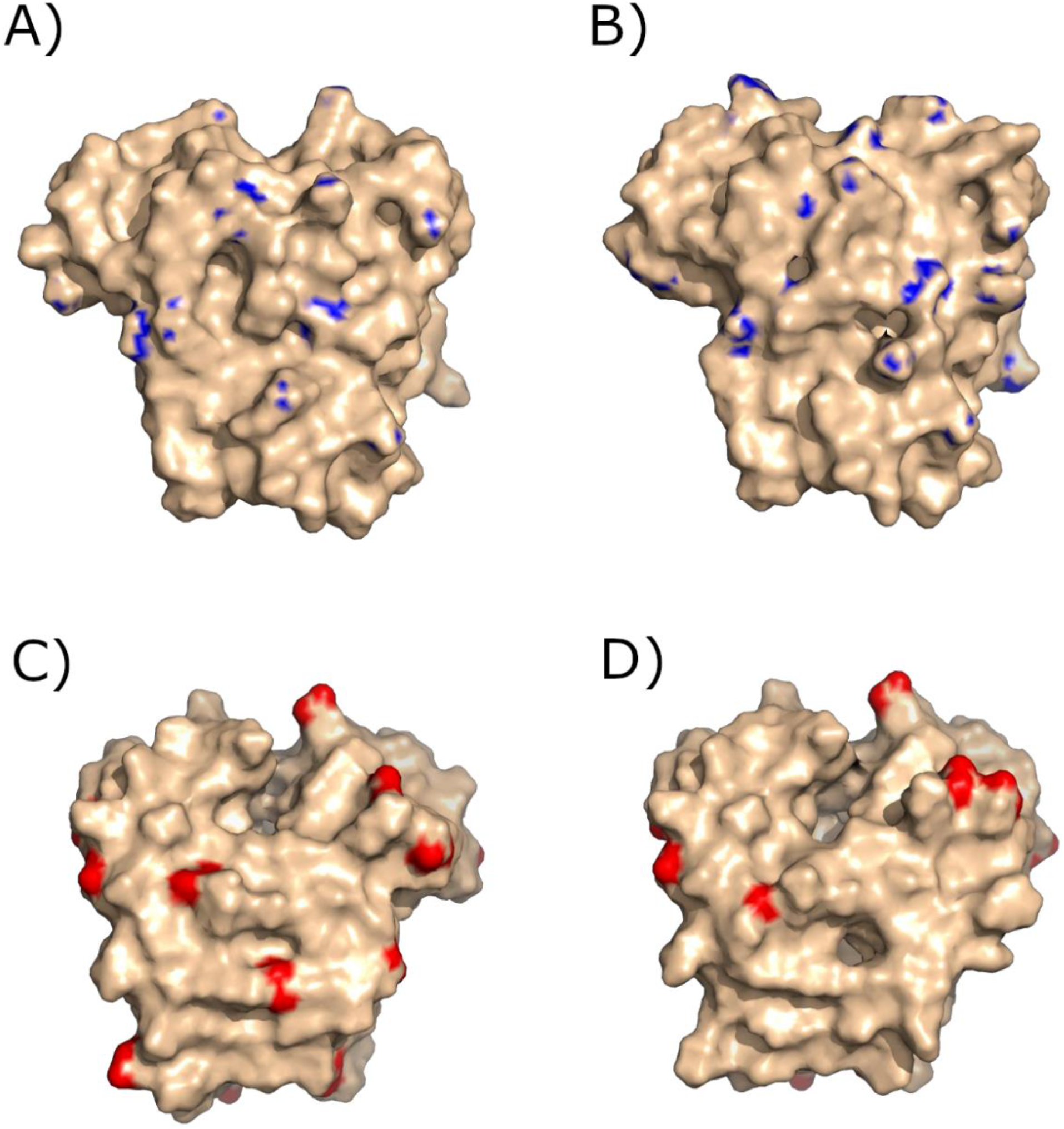
Contribution of positively charged atoms to solvent-accessible surface area for myoglobin models from **(A)** ostrich and **(B)** Emperor penguin, and contribution of negatively charged atoms to solvent-accessible surface area for myoglobin models from **(C)** mallard and **(D)** tufted duck.

### Positive selection in penguins and high purifying selection in galliforms

We investigated the intensity of evolutionary pressure on various myoglobins by calculating selective pressure (ω), or dN/dS ratios, for 68 of the 73 avian species in our dataset. Selection pressures for the following species could not be calculated due to sequence misalignment: tufted duck, North Island brown kiwi, saker falcon (*Falco cherrug*), and peregrine falcon (*Falco peregrinus*). The rest of the sequences were aligned with the common pigeon (*Columba livia*) myoglobin as the reference.

Our results show that the myoglobins of penguins have the largest selection pressure ratios (ω), which were extremely high compared to all other avian myoglobins (figure 4B). Although the selective pressures of all species were below 1, the relatively high selective pressures of the Adélie penguin and Emperor penguin, which were the only penguins in our dataset, might indicate intense positive selective pressure in the recent evolutionary past. Together with findings of highest positively charged solvent-accessible surface area, lowest negatively charged solvent-accessible surface area, and relatively high net charge, the finding of highest selective pressures (ω) strongly suggests evolution via natural selection of unusually positive electrostatic properties in the myoglobins of penguins, most likely by the substitution of neutral or negatively charged amino acids with positively charged amino acids especially near the solvent-accessible surface. These biochemical properties would have provided penguins a fitness advantage by reducing the aggregation propensity of their myoglobins, allowing greater muscle oxygen storage (in the form of increased intracellular myoglobin concentration levels) for extended underwater foraging.

Results also show that galliforms have the lowest selective pressure ratios, which suggests relatively high purifying selection on the myoglobins of this order. With the exception of the sunbittern (*Eurypyga helias*), which has the 4^th^ lowest selective pressure, the six lowest selective pressure ratios (figure 4B) belonged to all the galliforms in our dataset: the common pheasant (*Phasianus colchicus*), which had the lowest selective pressure ratio, followed by turkey (*Meleagris gallopavo*), chicken (*Gallus gallus*), Japanese quail (*Coturnix japonica*), and helmeted guineafowl (*Numida meleagris*). All of these are domesticated species, with the exception of the common pheasant, although a recent study found evidence of partial domestication of the pheasant in one of the earliest Neolithic agricultural sites.^33^ Perhaps myoglobin function is critical for the appearance of health and hence selective breeding success in birds undergoing domestication, which could explain the unusually high purifying selection that seems to be operating on the myoglobins of these birds. The exact nature of the relationship between myoglobin function and galliform biology should be investigated by future studies.

### High myoglobin net charge in some non-diving avian species

We found unusual myoglobin electrostatic properties in several non-diving avian species: kiwis, golden eagles, pelicans, and geese. The North Island brown kiwi myoglobin had the second highest predicted net charge (+5.31*e*, figure 4A) and among the lowest negatively charged atomic contributions to solvent-accessible surface area (figure 4D). The other kiwi species in our dataset, the Okarito kiwi (*Apteryx rowi*), had the fourth highest positively charged atomic contribution to solvent-accessible surface area (figure 4C) and a lower-than-average negatively charged atomic contribution to solvent-accessible surface area (figure 4D). We do not know the function of such high net charge in kiwis, as kiwis are not known to have an aquatic evolutionary history. However, our data predicts that they might have above-average muscle myoglobin concentration levels, which would indicate environmental hypoxia in their current or past environments.

The third most highly charged avian myoglobin (+4.09*e*) in the dataset belonged to the golden eagle, a high-altitude dwelling species. A previous study on golden eagle hemoglobin identified unique structural features that increase its oxygen affinity and hence allow the species to survive high-altitude hypoxia.^34^ The high net charge of golden eagle myoglobin (figure 4A) might be another molecular adaptation for hypoxic stresses at high altitudes. Given the established association between myoglobin net charge and muscle myoglobin concentration,^7^ and the fact that high-altitude vertebrates tend to possess higher concentrations of muscle myoglobin compared to their lowland counterparts,^35–37^ we predict that golden eagles possess unusually high concentrations of myoglobin in their muscles, which requires the high net charge for protection against aggregation. Experimental measurement of muscle myoglobin content in this species is needed to confirm this prediction.

One of the most highly charged myoglobins belonged to the Dalmatian pelican (*Pelecanus crispus*, +3.24*e*, figure 4A), which also had one of the highest contributions of positively charged atoms (figure 4C) and one of the lowest contributions of negatively charged atoms (figure 4D) to solvent-accessible surface area. Modern pelicans do not forage underwater for long periods of time, so we suspect their high net charge is a vestigial remnant of a more aquatic evolutionary past. If true, this would be similar to the case of modern elephants, which do not dive but have high myoglobin net charge due to their aquatic evolutionary history.^7^

### Inter-class differences in myoglobin net charge

Generally, we found that birds possess the most highly charged myoglobins while reptiles and amphibians possess the most negatively charged myoglobins (figure 7). The median and average myoglobin net charge for all vertebrates is +1.35*e* and +1.37*e*, respectively. Wilcoxon rank sum tests confirmed that the median net charge of bird myoglobin (+2.11*e*) was statistically significantly higher than that of mammalian myoglobin (+1.18*e*), reptilian myoglobin (−0.34*e*), and fish myoglobin (+1.41*e*) at the 5% significance level (p-values = 8.15×10^−7^, 2.37×10^−9^, and 0.02 for comparison of birds with mammals, reptiles, and fish, respectively), and that the median net charge of reptilian myoglobin (−0.34*e*) was statistically significantly lower than that of other classes (p-values = 5.45×10^−6^, 2.37×10^−9^, and 2.29×10^−4^ for comparison of reptiles with mammals, birds, and fish, respectively). Within reptiles, we noticed that snakes tended to have the most negatively charged myoglobins.

**Figure 7.**
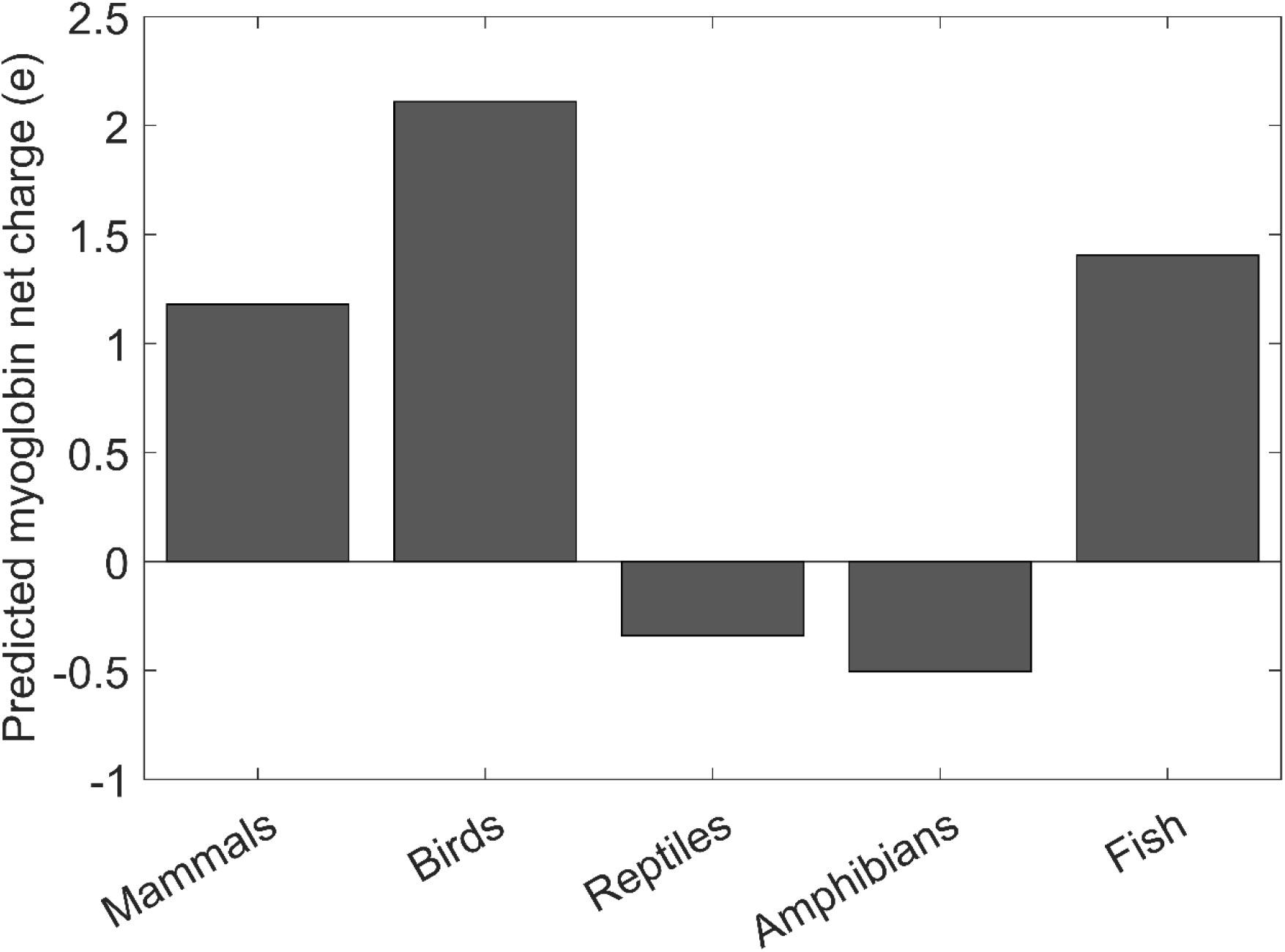
Predicted median myoglobin net charge in mammals, birds, reptiles, amphibians, and fish.

The inter-class differences in myoglobin net charge cannot be definitively attributed to differences in muscle myoglobin content, because studies of avian and reptilian muscle did not find myoglobin concentrations that are particularly different from those of mammals.^37–39^ The low myoglobin net charge of reptiles and amphibians (figure 7) might be related to their ectothermy, but an exact mechanism by which a higher myoglobin net charge would be beneficial for homeothermic vertebrates remains to be identified. Perhaps the lower myoglobin net charge of reptiles is a byproduct, or even a cause for, the lower oxygen affinities of reptilian globins compared to those of mammals.^38,40^

### Inverse function describes relationship between myoglobin concentration and net charge better than logarithmic function

We reexamined the mathematical relationship between experimentally measured myoglobin concentrations and myoglobin net charge with a larger dataset than previous studies.^7^ Specifically, our dataset includes myoglobin concentration data from the following five additional species which were not included in the analyses of previous studies: common dolphin (*Delphinus delphis*), Pacific white-sided dolphin (*Lagenorhynchus obliquidens*), beluga whale (*Delphinapterus leucas*), sea otter (*Enhydra lutris*), and North American beaver (*Castor canadensis*).^7^ We tested the fit of various mathematical functions, including logistic, parabolic, exponential (up to the 7^th^ degree polynomial), inverse, and logarithmic, which was the model proposed by previous studies. Although the inverse function did not have the best quantitative measures of fit (e.g. R^2^ and RMSE), it was visually the best function for describing the shape of the data. As seen in figure 8, the inverse function describes the relationship better than the logarithmic function, as the inverse function accounts for the horizontal asymptote in the data and the much more rapid decline of net charge at lower concentrations.

**Figure 8.**
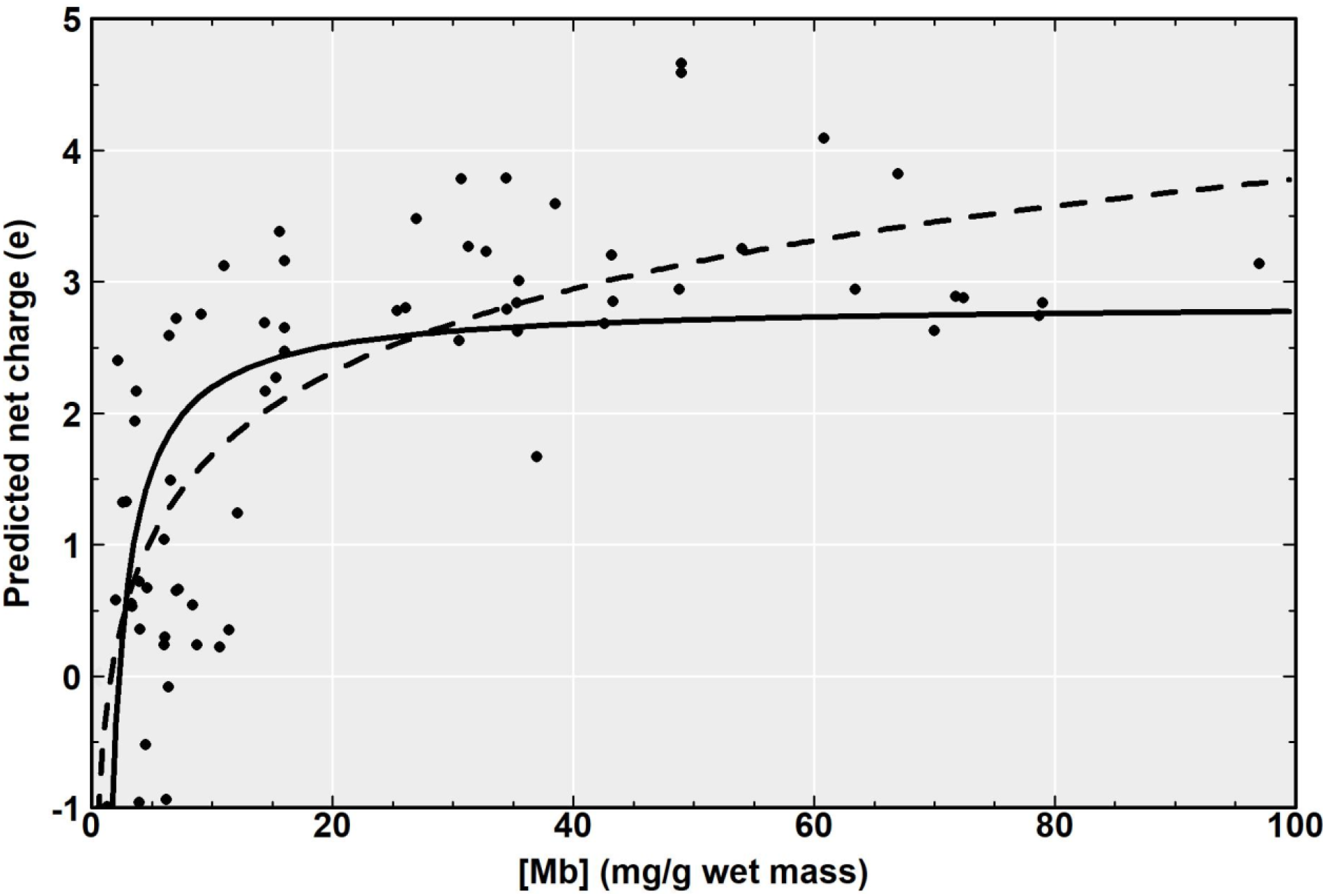
Predicted myoglobin net charge and experimentally determined muscle myoglobin concentrations in mammals. Dashed line represents logarithmic regression and solid line represents the inverse function.

Furthermore, we attempt to provide physical explanations for the terms in the inverse equation, which previous studies did not do. We believe the relationship can be described in terms of a physical contribution (horizontal asymptote, represented by *a*) and biological fitness contribution (rapid decline of net charge at low concentrations, represented by *b*):

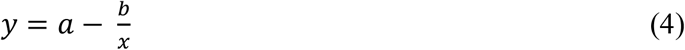

where *a* and *b* are empirically determined constants: 2.837 and 6.382, respectively. As seen in figure 8, the relationship between predicted net charge and myoglobin concentration plateaus around a horizontal asymptote of +2.837*e* in net charge, corresponding to greater than ∼40 mg/g myoglobin concentration levels. In other words, once myoglobin intracellular concentration reaches this threshold, a net charge of ∼+2.837*e* is needed to prevent myoglobin aggregation. Using Coulomb’s law, we can calculate the force required to keep two myoglobin proteins from aggregating upon physical contact with each other at the speeds they move within the cell, where *r =* 3.5 × 10^−9^ meters is the diameter of myoglobin^41^:

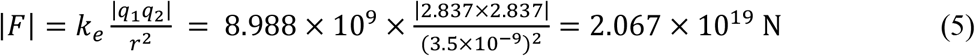

In the absence of this physical constraint, when myoglobin concentration levels are low and there is little risk of aggregation, unknown biological fitness constraints keep the myoglobin net charge closer to neutral.

### High myoglobin net charge is negatively associated with herbivory

The macroevolutionary transition of cetaceans from terrestrial herbivores to fully aquatic myoglobin-rich carnivores prompted us to investigate whether any relationships exist between myoglobin net charge and mammal diet. Due to relative lack of carnivorous species in our dataset, we compared the predicted net charge of herbivores and omnivores only. We found that the median net charge of omnivores (+1.03*e*) was double that of herbivores (+0.37*e*), and a Wilcoxon rank sum test confirmed that this difference was statistically significant (p-value = 0.0023). The causal relationship, if any, between diet and myoglobin net charge remains to be identified.

## Conclusion

Although myoglobin is one of the most well-studied proteins, its biochemical properties and evolution in non-mammalian vertebrates remains poorly characterized. Here, we used a structural bioinformatics approach to investigate the evolution of myoglobin in 302 species from a variety of vertebrate classes, and calculated selection pressures for 68 avian species. We found that penguins had the highest selection pressure ratios, and also the highest contribution of positively charged atoms to solvent-accessible surface area, which suggests they evolved supercharged myoglobins to prevent aggregation at the high intracellular concentrations required for long-duration underwater hypoxia. Positively charged atoms on the solvent-accessible surface of myoglobin were more indicative of high myoglobin content (and hence aggregation resistance) than net charge. We also found the lowest selection pressures in galliform myoglobins, indicating relatively high purifying selection in these species. Finally, we found inter-class differences in net charge, and formulated an equation that describes the relationship between myoglobin net charge and concentration better than the previously suggested logarithmic function.

One limitation of this study is that few non-mammalian data could be used to derive the current model in Equation 4, as there are only two amphibians among the deposited sequences in online databases, and concentration data for non-mammalian vertebrates was scarce. We expect that a more accurate model can be derived when more non-mammalian data become available.

## Acknowledgments

We thank Xiaoqiang Huang for feedback and stimulating discussions. This work used the Extreme Science and Engineering Discovery Environment (XSEDE), which is supported by the National Science Foundation grant ACI1548562. This work was supported in part by National Institutes of Health (GM083107, GM136422, AI134678, OD026825 to Y.Z.), and the National Science Foundation (DBI1564756 and IIS1901191 to Y.Z. and MTM2025426 to Y.Z.).

## Data Availability

The data underlying this article are available in the article and in its online supplementary material.

## Notes

### Competing Interest Statement

The authors have declared no competing interest.

### Summary of Updates

Sections on hydrophobicity have been removed and new findings related to selection pressures and associations between net charge and herbivory have been added.

## References

1. Gros G, Wittenberg BA, Jue T. Myoglobin’s old and new clothes: From molecular structure to function in living cells. J Exp Biol. 2010;213:2713–2725. doi:10.1242/jeb.043075

2. Lin Y, Wang J, Lu Y. Functional tuning and expanding of myoglobin by rational protein design. Sci China Chem. 2014;57(3):346–355. doi:10.1007/s11426-014-5063-5

3. Van Der Meer DLM, Van Den Thillart GEEJM, Witte F, et al. Gene expression profiling of the long-term adaptive response to hypoxia in the gills of adult zebrafish. Am J Physiol - Regul Integr Comp Physiol. 2005;289(5):R1512–R1519. doi:10.1152/ajpregu.00089.2005

4. Ferreras JM, Ragucci S, Citores L, Iglesias R, Pedone P V., Di Maro A. Insight into the phylogenetic relationship and structural features of vertebrate myoglobin family. Int J Biol Macromol. 2016;93:1041–1050. doi:10.1016/j.ijbiomac.2016.09.065

5. Wright TJ, Davis RW. Myoglobin oxygen affinity in aquatic and terrestrial birds and mammals. J Exp Biol. 2015;218:2180–2189. doi:10.1242/jeb.119321

6. Berenbrink M. The role of myoglobin in the evolution of mammalian diving capacity – The August Krogh principle applied in molecular and evolutionary physiology. Comp Biochem Physiol Part A Mol Integr Physiol. 2021;252:110843. doi:10.1016/J.CBPA.2020.110843

7. Mirceta S, Signore A V., Burns JM, Cossins AR, Campbell KL, Berenbrink M. Evolution of mammalian diving capacity traced by myoglobin net surface charge. Science (80-). 2013;340:1234192. doi:10.1126/science.1234192

8. Serebryany E, Woodard JC, Adkar B V., Shabab M, King JA, Shakhnovich EI. An Internal Disulfide Locks a Misfolded Aggregation-prone Intermediate in Cataract-linked Mutants of Human γD-Crystallin. J Biol Chem. 2016;291(36):19172. doi:10.1074/JBC.M116.735977

9. Aguzzi A, O’Connor T. Protein aggregation diseases: Pathogenicity and therapeutic perspectives. Nat Rev Drug Discov. 2010;9(3):237–248. doi:10.1038/nrd3050

10. Cleland JL, Powell MF, Shire SJ. The development of stable protein formulations: A close look at protein aggregation, deamidation, and oxidation. Crit Rev Ther Drug Carrier Syst. 1993;10(4):307–377.

11. Mitragotri S, Burke PA, Langer R. Overcoming the challenges in administering biopharmaceuticals: Formulation and delivery strategies. Nat Rev Drug Discov. 2014. doi:10.1038/nrd4363

12. Olsson MHM, Søndergaard CR, Rostkowski M, Jensen JH. PROPKA3: Consistent Treatment of Internal and Surface Residues in Empirical p<italic>K</italic>_a_Predictions BT -Journal of Chemical Theory and Computation. J Chem Theory Comput. 2011;7(2):525–537.

13. Li Y, Zheng W, Zhang C, et al. Protein 3D Structure Prediction by D-I-TASSER in CASP14. In: CASP 14 Abstract Book. ; 2020:339–341.

14. Yang J, Yan R, Roy A, Xu D, Poisson J, Zhang Y. The I-TASSER suite: Protein structure and function prediction. Nat Methods. 2015;12(1):7–8. doi:10.1038/nmeth.3213

15. Zheng W, Li Y, Zhang C, et al. Protein structure prediction using deep learning distance and hydrogen-bonding restraints in CASP14. Proteins Struct Funct Bioinforma. 2021;89(12):1734–1751. doi:10.1002/PROT.26193

16. Zhang C, Zheng W, Mortuza SM, Li Y, Zhang Y. DeepMSA: Constructing deep multiple sequence alignment to improve contact prediction and fold-recognition for distant-homology proteins. Bioinformatics. 2020;36:2105–2112. doi:10.1093/bioinformatics/btz863

17. Li Y, Zhang C, Zheng W, et al. Protein inter-residue contact and distance prediction by coupling complementary coevolution features with deep residual networks in CASP14. Proteins. 2021;89(12):1911–1921. doi:10.1002/PROT.26211

18. Zhang Y, Skolnick J. SPICKER: A clustering approach to identify near-native protein folds. J Comput Chem. 2004;25(6):865–871. doi:10.1002/jcc.20011

19. Zhang J, Liang Y, Zhang Y. Atomic-level protein structure refinement using fragment-guided molecular dynamics conformation sampling. Structure. 2011;19(12):1784–1795. doi:10.1016/j.str.2011.09.022

20. Huang X, Pearce R, Zhang Y. FASPR: An open-source tool for fast and accurate protein side-chain packing. Bioinformatics. 2020;36(12):3758–3765. doi:10.1093/bioinformatics/btaa234

21. Zhang Y, Skolnick J. Scoring function for automated assessment of protein structure template quality. Proteins Struct Funct Genet. 2004;57:702–710. doi:10.1002/prot.20264

22. Xu J, Zhang Y. How significant is a protein structure similarity with TM-score = 0.5? Bioinformatics. 2010;26(7):889–895. doi:10.1093/bioinformatics/btq066

23. Zhang Y. I-TASSER server for protein 3D structure prediction. BMC Bioinformatics. 2008;9(1):40. doi:10.1186/1471-2105-9-40

24. Frishman D, Argos P. Knowledge-based protein secondary structure assignment. Proteins Struct Funct Bioinforma. 1995;23:566–579. doi:10.1002/prot.340230412

25. Baker NA, Sept D, Joseph S, Holst MJ, McCammon JA. Electrostatics of nanosystems: Application to microtubules and the ribosome. Proc Natl Acad Sci U S A. 2001;98(18):10037–10041. doi:10.1073/pnas.181342398

26. Prum RO, Berv JS, Dornburg A, et al. A comprehensive phylogeny of birds (Aves) using targeted next-generation DNA sequencing. Nature. 2015;526:569–573. doi:10.1038/nature15697

27. Kissling WD, Dalby L, Fløjgaard C, et al. Establishing macroecological trait datasets: digitalization, extrapolation, and validation of diet preferences in terrestrial mammals worldwide. Ecol Evol. 2014;4(14):2913–2930. doi:10.1002/ECE3.1136

28. Zhang C, Zheng W, Cheng M, Omenn GS, Freddolino PL, Zhang Y. Functions of Essential Genes and a Scale-Free Protein Interaction Network Revealed by Structure-Based Function and Interaction Prediction for a Minimal Genome. J Proteome Res. 2021. doi:10.1021/acs.jproteome.0c00359

29. Ponganis PJ, Costello ML, Starke LN, Mathieu-Costello O, Kooyman GL. Structural and biochemical characteristics of locomotory muscles of emperor penguins, Aptenodytes forsteri. Respir Physiol. 1997;109(1):73–80. doi:10.1016/S0034-5687(97)84031-5

30. Stephenson R, Turner DL, Butler PJ. The Relationship Between Diving Activity and Oxygen Storage Capacity in the Tufted Duck (Aythya Fuligula). J Exp Biol. 1989;141:265–275.

31. Weber RE, Hemmingsen EA, Johansen K. Functional and biochemical studies of penguin myoglobin. Comp Biochem Physiol -- Part B Biochem. 1974;49(2):197–204. doi:10.1016/0305-0491(74)90154-0

32. Kagen LJ, Linder S. Immunological Studies of Duck Myoglobin. Proc Soc Exp Biol Med. 1968;128(2):438–441. doi:10.3181/00379727-128-33032

33. Barton L, Bingham B, Sankaranarayanan K, Monroe C, Thomas A, Kemp BM. The earliest farmers of northwest China exploited grain-fed pheasants not chickens. Sci Reports 2020 101. 2020;10(1):1–7. doi:10.1038/s41598-020-59316-5

34. Ali A, Rehman T, Roshan A. Analysis of Oxygen Affinity of the Major Hemoglobin Component HbA from Aquila chrysaetos (Golden Eagle). J Nat Sci Res. 2014;4(9):55–61.

35. Dawson NJ, Ivy CM, Alza L, et al. Mitochondrial physiology in the skeletal and cardiac muscles is altered in torrent ducks, Merganetta armata, from high altitudes in the Andes. J Exp Biol. 2016;219(23):3719–3728. doi:10.1242/jeb.142711

36. Vaughan BE, Pace N. Changes in myoglobin content of the high altitude acclimatized rat. Am J Physiol. 1956;185(3):549–556. doi:10.1152/ajplegacy.1956.185.3.549

37. Xin Y, Tang X, Wang H, et al. Functional characterization and expression analysis of myoglobin in high-altitude lizard Phrynocephalus erythrurus. Comp Biochem Physiol Part - B Biochem Mol Biol. 2015;188:31–36. doi:10.1016/j.cbpb.2015.06.004

38. Weber RE, Johansen K, Abe AS. Myoglobin from the burrowing reptile Amphisbaena alba. Concentrations and functional characteristics. Comp Biochem Physiol -- Part A Physiol. 1981;68(2):159–165. doi:10.1016/0300-9629(81)90336-4

39. Pages T, Planas J. Muscle myoglobin and flying habits in birds. Comp Biochem Physiol -- Part A Physiol. 1983;74(2):289–294. doi:10.1016/0300-9629(83)90602-3

40. Pough FH. Blood oxygen transport and delivery in reptiles. Integr Comp Biol. 1980;20(1):173–185. doi:10.1093/icb/20.1.173

41. Papadopoulos S, Jürgens KD, Gros G. Protein diffusion in living skeletal muscle fibers: dependence on protein size, fiber type, and contraction. Biophys J. 2000;79(4):2084. doi:10.1016/S0006-3495(00)76456-3

